# Spontaneous but not voluntary eye blinks during spatial working memory are associated with successful performance

**DOI:** 10.1101/345306

**Authors:** Andrew S. Lee, Baaba K. Blankson, Peter Manza, Jonathan F. O’Rawe, Craig Evinger, Hoi-Chung Leung

## Abstract

Spontaneous eye blink rate (SBR) has been associated with central dopamine (DA) levels, raising the intriguing possibility that SBR is related to cognitive functions dependent on DA, such as spatial working memory (WM). We tested this hypothesis in two behavioral experiments, examining the relationship between SBR, WM load and individual differences in spatial WM performance in 126 young adults. In Experiment 1, we examined the temporal profile of SBR during a spatial delayed recognition task requiring maintenance of 1, 2, 4, 6 or 7 dot locations. We observed a suppression in SBR during dot- and recognition probe-presentation, and a significant increase in SBR afterwards. High performers showed significantly lower SBR than low performers during the first 500 ms of the delay period. In Experiment 2, we used a similar spatial WM task as Experiment 1 to test whether an instructed voluntary blink during the early delay would directly dampen WM performance. While the temporal dynamics of SBR across task events were comparable to Experiment 1, WM performance was not significantly different between the voluntary blink and no blink conditions. Together, these results suggest that spontaneous but not voluntary eye blinking is closely linked to spatial WM, and that lower SBR during WM encoding and early phase of maintenance is associated with better WM task performance.

## INTRODUCTION

Spontaneous blinking is a critical process that allows maintenance of corneal tear film (Sibony and Evinger, 1998; Evinger, 2010), but the number of spontaneous blinks varies depending on the individuals’ behavioral state (Doughty, 2001). The variability in frequency and number of blinks does not account for any immediate physiological needs (Al-Abdulmunem, 1999), suggesting that eye blinking is modulated by cognitive demands that are constantly imposed on an individual (Stern, Walrath, & Goldstein, 1984). In fact, the average spontaneous blink rate (SBR) is lowest during reading and highest during conversation (Ponder and Kennedy, 1927; Karson et al., 1981; Fogarty and Stern, 1989; Orchard and Stern, 1991; Doughty, 2001; Pivik and Dykman, 2004). This variation may reflect a balance between blinking to satisfy immediate physiological needs and suppressing eye blinks during visual processing (Bristow et al., 2005), as it is crucial to minimize interference or visual information loss as a consequence of eye blinking. However, the temporal pattern of SBR during cognitive processing and its implications for task performance remains unclear.

While the timing of spontaneous blinks exhibits a consistent temporal organization under no explicit tasks (Kaminer et al., 2011), it changes across different temporal events during cognitive task performance (Stern, Walrath, & Goldstein, 1984; Hoppe, Helfmann, & Rothkopf, 2018). Many studies showed that SBR decreases in anticipation of and during stimulus presentation and increased or rebound after stimulus offset (Baumstimler & Parrot, 1971; Stern, Walrath, & Goldstein, 1984; Bauer, Goldstein, & Stern, 1987; Goldstein, Bauer, & Stern, 1992; Ohira, 1996; Boehm-Davis, Gray, & Schoelles, 2000; Fukuda, 1994; Fukuda, 2001; Ichikawa & Ohira, 2004; Nakano et al., 2009; Oh, Jeong, & Jeong, 2012). A lower SBR has been observed in high cognitive load conditions compared to low cognitive load conditions in variety of tasks including a bomber mission simulation (Stern & Skelly, 1984), air traffic control game (Brookings, Wilson, Swain, 1996), Sternberg memory task (Goldstein, Bauer, & Stern, 1992), semantic priming (Ohira, 1996), flight simulation (Veltman & Gaillard, 1998), Multi-Attribute Test Battery (Fournier, Wilson, & Swain, 1999), visual tracking (Van Orden, Jung, & Makeig, 2000), anti-air-warfare simulation (Van Orden, Limbert, Makeig, & Jung, 2001), lexical decision making (Ichikawa and Ohira, 2004), digit sorting (Siegle, Ichikawa, & Steinhauer, 2008). Together, these prior studies suggest that SBR suppression pattern is not random, but critical for stimulus encoding and task performance. These earlier studies thus suggest that SBR is potentially indicative of a specific point in which information processing is occurring, therefore, the temporal dynamics of SBR can be used to index the quality and type of cognitive processing taking place.

Aside from being a noninvasive, inexpensive, and easy-to-obtain physiological measure of a person’s cognitive state, SBR is suggested to be an indirect measure of central dopamine (DA) levels (see review by Jongkees and Colzato, 2016). Studies using pharmacological manipulations showed that treatment of DA receptor 1 (D_1_R) agonists in rodents and non-human primates increases SBR while antagonists decrease SBR (Desai et al., 2007; Kotani et al., 2016; Karson, Staub, Kleinman, & Wyatt, 1981; Elsworth et al., 1991; Lawrence and Redmond, 1991; Jutkiewicz and Bergman, 2004; Kleven and Koek, 1996; Taylor et al., 1999; Kaminer et al., 2011). More recently, a more comprehensive study showed that PET measure of striatal D_2_R availability and *in vivo* measure of striatal DA receptor density is closely related to individual differences in SBR across adult male vervet monkeys (Groman et al., 2014). Although the relationship between SBR and specific DA receptor occupancy is less investigated in humans and remains controversial (Dang et al., 2017; Sescousse et al., 2018), studies of DA-related neurological disorders such as Parkinson’s disease indicate that central DA activity also modulates SBR in humans (Karson, LeWitt, Calne, & Wyatt, 1982; Golbe, Lepre, & Davis, 1989; Deuschl and Goddemeier, 1998; Korošec et al., 2006).

One particular cognitive process tightly coupled to central DA activity is spatial working memory (WM), a fundamental cognitive process that supports maintenance and manipulation of information necessary for completing complex tasks. Rodent and non-human primate models showed that D_1_R in the prefrontal cortex play an essential role in spatial WM maintenance (Arnsten, Cai, Murphy, Goldman-Rakic, 1994; Cai and Arnsten, 1997; Castner and Goldman-Rakic, 2004; Sawaguchi and Goldman-Rakic, 1991, 1994; Goldman-Rakic, 1992; Robbins, 2000). Studies of humans have demonstrated that peripheral administration of D_2_R agonist or antagonists can influence spatial WM capacity and highlighted the role of D_2_R activity in WM (Luciana, Depue, Arbisi, & Leon, 1992; Luciana and Collins, 1997; Kimberg, D’Esposito, & Farah, 1997; Luciana, Collins, & Depue, 1998; Kimberg, Aguirre, Lease, & D’Esposito, 2001; Kimberg & D’Esposito, 2003). Other studies in humans using PET imaging have shown that striatal DA synthesis capacity is positively correlated with WM capacity of a listening span task (Gibbs & D’Esposito, 2005; Cools et al., 2008; Landau et al., 2009). However, our knowledge of the specific contributions of DA in WM in human remains limited due to the challenges of expensive pharmacological imaging. While some studies have examined the relationship between SBR and WM performance (Irwin, 2014; Dreisbach et al., 2005; Tharp and Pickering, 2011; Zhang et al., 2015; Rac-Lubashevsky, Slagter, & Kessler, 2017), they used SBR measured during resting conditions as a proxy of central DA system and not during different phases of WM. Therefore, individual differences in eye blink activity during WM processing and task performance is unclear.

### The Current Study

While the temporal dynamics of SBR have been studied using a number of cognitive tasks, these tasks often probe multiple cognitive processes simultaneously. To distinguish the pattern of SBR across different cognitive processes, we utilized a spatial WM paradigm, which includes temporally separated stages of cognitive processing (stimulus encoding, maintenance, and recognition), and manipulated the WM load. This paradigm also allows us to examine the effects of cognitive load on SBR after stimulus display without the anticipatory blink-rebound effects from priming cues. To characterize individual differences in SBR associated with WM capacity, we computed individual K-score and compared SBR differences between participants of high versus low K-score. We hypothesized that blink suppression during critical periods of a spatial WM task is associated with successful task performance. Given that both SBR and spatial WM are associated with the central DA system, examining SBR during spatial WM may provide more temporally precise data to complement prior studies on spatial WM and the central DA system.

### Overview of Experiments

In two behavioral experiments, participants completed a spatial delayed recognition paradigm while eye blink activity was monitored and recorded. In Experiment 1, we recorded eye blink activity using vertical electrooculogram (VEOG) to characterize the temporal pattern of SBR during different events of spatial WM. We replicated previous findings such that SBR was suppressed during target and probe display and followed by a significant rebound. Participants with higher K-score showed significantly lower SBR during the first 500 ms of the delay period than participants of lower k-score, and this difference was WM load dependent.

In Experiment 2, we used a similar paradigm as Experiment 1 (but a shorter stimulus display), and measured eye blink activity using an eye tracker. Eye blink activity was recorded while performing the WM task performance and watching a video of natural scenes (eye blink activity in a condition without an explicit task). To test whether WM performance is influenced by eye blinking per se, we instructed the participants to either blink or not blink during the first 500ms of the delay period. We expected that a voluntary blink during this critical period would impair WM performance compared to participants instructed *not* to blink. While the temporal dynamics of SBR across task events were comparable to Experiment 1, WM performance was not significantly different between the blink and no blink groups. Also, there was a significant positive correlation with SBR measured while participants watched a video of natural scenes and WM capacity.

## METHODS

### Experiment 1

#### Participants

Sixty healthy participants (34 females, mean age = 20.8 ± 3 [range: 18 to 39], all right handed) were recruited via Stony Brook University’s Psychology subject pool, consented and received course credit for their participation. Prior to the experiment, participants completed a consent form and a handedness form according to the Edinburgh Inventory (Oldfield, 1971). They all had normal or corrected-to-normal vision and self-reported no history of psychiatric and/or neurological disorders. The study protocol was approved by the local Institutional Review Board.

#### General Procedures, Apparatus and Stimuli

The spatial WM span task for the present analysis was part of a larger study, which included event related potentials (ERP) recording during the completion of a more complex spatial WM task. Participants performed two practice blocks of ten trials each before the actual spatial WM span task. The actual task lasted approximately 7.5 minutes and consisted of eighty trials in total with a 10-second break after every twenty trials. Testing was conducted in a sound-attenuated room with dim lighting. A personal computer (PC) with the Windows XP operating system (Pentium 4, Dell Dimension 5100; Dell Computer Company, Round Rock, TX) was used to deliver visual stimuli and record button presses. The monitor display mode was set to 1024 × 768 pixels, 100 Hz refresh rate, and 32-bit color. The monitor screen was positioned in the center of participants’ visual field. Visual presentation was controlled, and response data was recorded with E-prime (Psychology Software Tools, Pittsburgh, PA). Visual stimuli were presented within an 8 × 8 grid (gray in color, RGB: 220,220,220) that extended over a visual angle of 12.7° × 12.7° against a white background. Dark gray dots were used as target stimuli and unfilled circles as probe stimuli. Both the target dots and probe circles had diameters of 1.59° in visual angle at a 3-foot viewing distance.

#### Spatial Working Memory Task

The spatial WM span task was a delayed recognition paradigm (Figure 1a). Each trial began with a 500 ms fixation cross followed by the target stimulus presentation for 500 ms. Participants were instructed to memorize the location of each dot presented on the grid. Afterwards, there was a delay of 2000 ms followed by a probe stimulus presented for 500 ms. The probe could either match one of the study stimuli locations or in a new location. Participants were instructed to make a response about whether or not the probe was in one of the remembered locations during the 1500 ms inter-trial interval (ITI). Each trial lasted 5000 ms, the target and probe dots were presented pseudorandomly in different locations. To reduce interference, dot locations were not reused in trials after they had been presented. Each participant completed total of 80 trials, with an equal amount of “yes” and “no” trials. WM load was manipulated as the number of dots presented in the memory set (1, 2, 4, 6 or 7 dots, 16 trials each).

**Figure 1.**
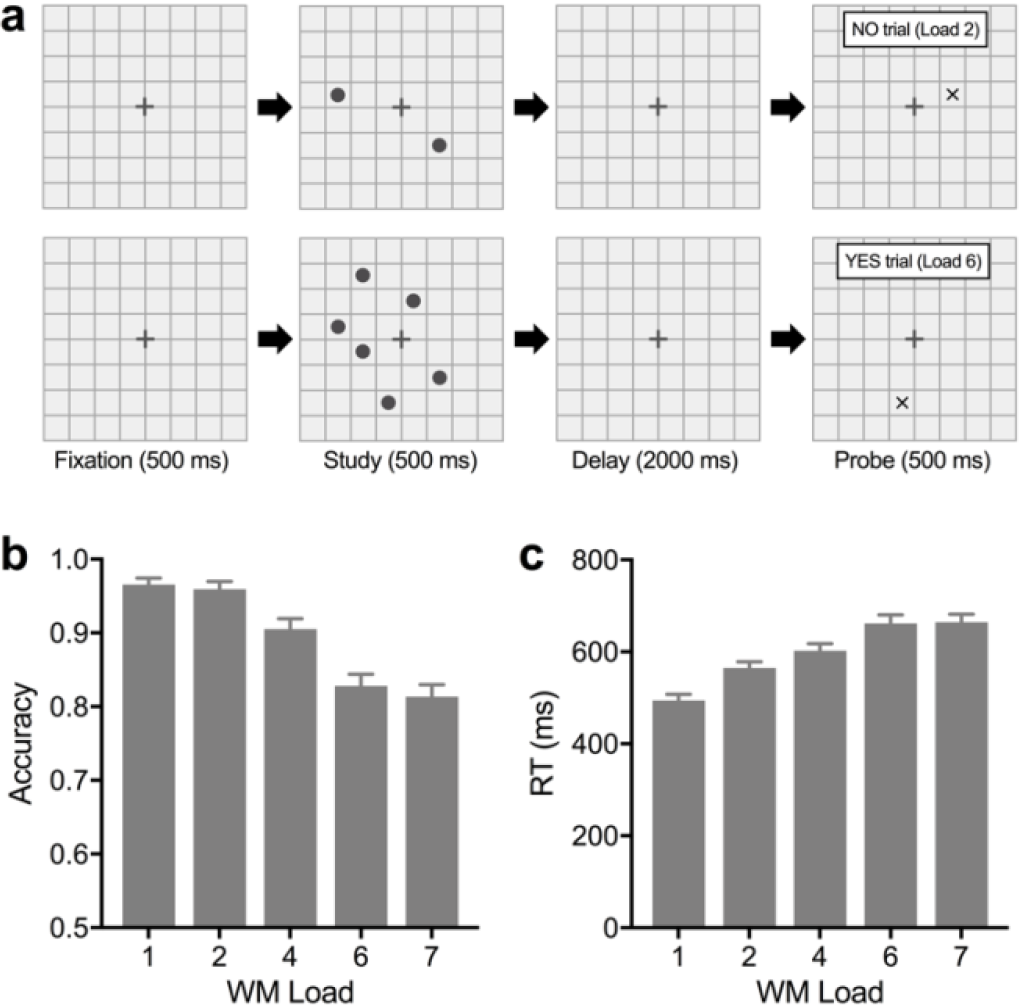
Schematic diagram of spatial WM task and behavioral results for Experiment 1. (a) The WM load consisted of 1, 2, 4, 6, or 7 dots presented in different locations on each trial, followed by a delay. After the delay, a probe appeared prompting a recognition response (yes or no). (b) accuracy (c) RT by WM load. Data are expressed as mean ± 1 SEM.

#### Data Analysis

Eye blink data and WM data were tested using SPSS Statistics 24. Accuracy and reaction time (RT) of different WM loads were analyzed with a repeated-measures ANOVA. We also calculated the K-score, which is an estimation of the number of items an individual can simultaneously hold in their WM (Cowan, 2001; Pashler, 1988). K-scores were calculated as K = N × (Hit Rate - False Alarm) where N is the WM load or number of to-be-remembered locations (Cowan, 2001; Pashler, 1988). Each participant’s WM capacity was calculated as an average of their K-scores from load 4, 6 and 7 (average K-score).

For the spontaneous eye blink analysis, we used data from the VEOG channel. A laboratory developed program identified blinks by eyelid position, velocity, and the occurrence of eye blinks for each participant. We first segmented the data into 5 s epochs (which is the total duration of each trial). Each epoch was baseline-corrected based on the average amplitude of the 100 ms pre-stimulus time bin. Blinks on incorrect trials were not separately analyzed since the total number of incorrect trials were too few, 6% on average of the 80 trials across all participants. For each participant and condition of interest (each memory load or correct trials), SBR was calculated by dividing the number of blinks during the specific condition by the total number of trials for that condition. We also examined temporal profile of SBR for each condition by dividing each trial into 500 ms bins and calculating SBR for each time bin (see Figure 2a).

**Figure 2.**
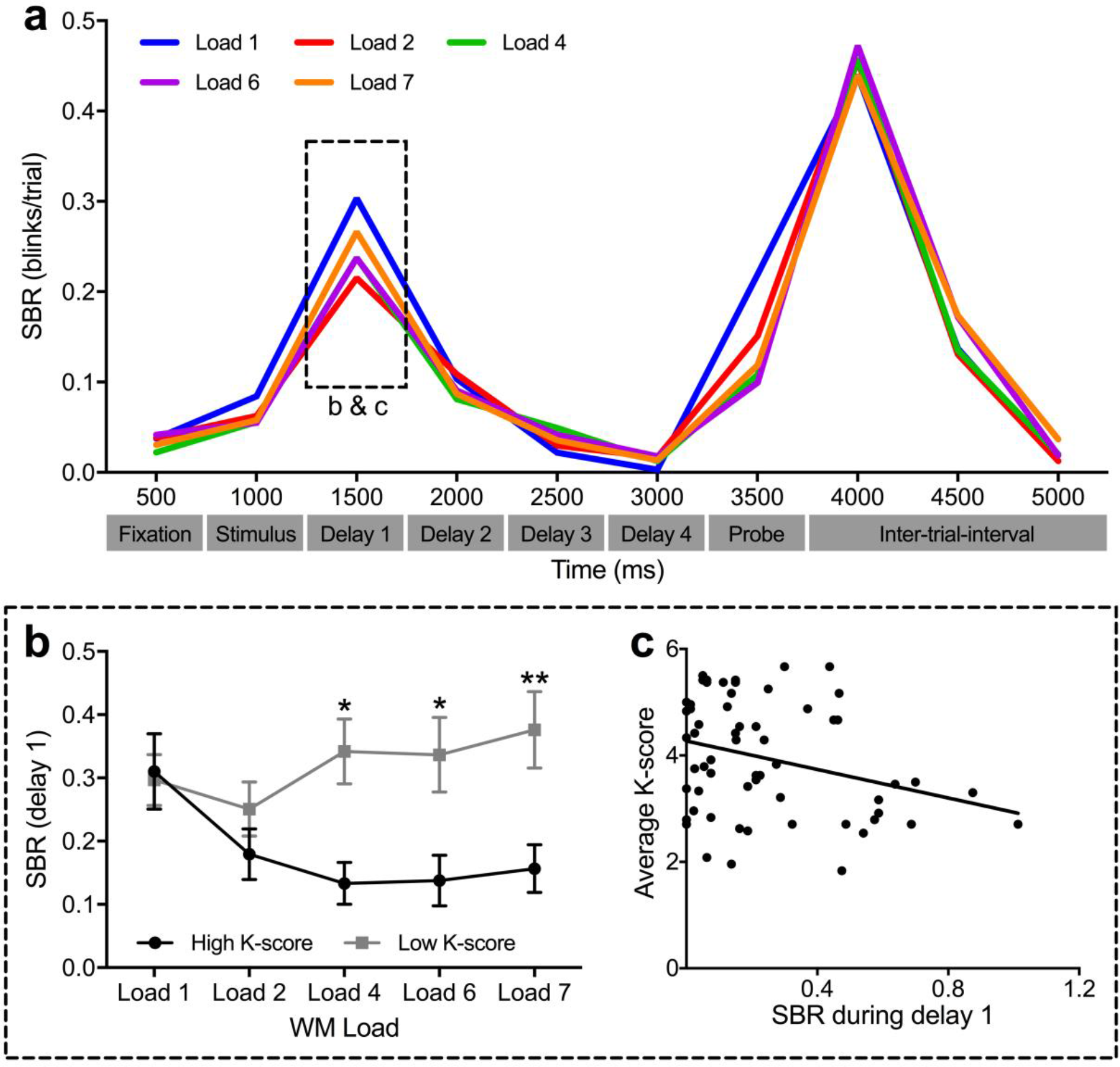
Time course of SBR during a full trial and relationship between SBR and WM capacity for Experiment 1. (a) Average SBR across 500 ms time bins for each WM load condition. (b) Differences in SBR during first 500 ms of the delay period (delay 1) between high K-score and low K-score performers by WM load. Data are expressed as mean ± 1 SEM. *Post hoc* comparison (Bonferroni correction); \**p* < .05 and \*\**p* < .01. (c) Correlation between SBR during time bin 1500 ms (“delay 1,” dotted-line box in (a)) and average K-score; the average of load 4, 6 and 7).

ANOVA analyses that violated the assumption of sphericity as indicated by Mauchly’s test, Greenhouse-Geisser corrected values were reported. A *post hoc* comparisons with Bonferroni corrections (significance was assumed at a value of *p* < .05) were used to explore any significant effects further. Correlations of SBR with performance measures were examined using a Pearson correlation, and significant correlations were plotted with a linear fit. To test any outliers, we calculated Cook’s distance, and excluded subjects that were above the cutoff (4/*n*, where *n* is number of participants). None of the subjects were outliers, therefore all subjects were included in the final analyses.

### Experiment 2

#### Participants

Another sixty-six healthy participants were recruited via Stony Brook University’s Psychology subject pool, consented and received course credit for their participation. They all had normal or corrected-to-normal vision and self-reported no history of psychiatric and/or neurological disorders. Due to technical difficulties of the eye tracker, participant compliance, and behavioral performance outlier (2 SD away from the mean), the final analysis included 49 participants (23 females, mean age = 19.2 ± 1.93 [range: 17 to 25], all right handed except one). The study protocol was approved by the local Institutional Review Board.

#### General Procedures, Apparatus and Stimuli

Participants performed a three-part experiment. The first part was to collect baseline SBR while participants watched a video. Participants then practiced one block of ten trials of the spatial WM task to be performed in the second (baseline WM) and third (WM with blink instructions) part of the experiment. The second and third parts were the actual spatial WM span task. Testing conditions were similar to Experiment 1, except the monitor display mode was set to 1680 × 1050 pixels, 100 Hz refresh rate, and 32-bit color, and a chinrest was used to control the distance between participants’ eyes and the computer monitor (3 feet). Visual stimuli were again presented within an 8 × 8 grid that extends a visual angle of 10.21° × 10.14° while the target dots and probe circles had diameters of 0.720° × 0.715° in visual angle. Participants were randomly assigned to conditions in which the left key of the joystick was either “yes” or “no” to control for handedness. Eye blinks and eye position were monitored and recorded using Eyelink 1000 (SR Research Ltd., Ontario, Canada).

#### Baseline Spontaneous Eye Blink Rate Measured While Watching Videos

Before the WM task, participants were shown a 10-minute video of natural scenes (1 scene/min) while their eye gaze and blinks were monitored and recorded. To reduce eye blinks evoked by gaze shifts, subjects were asked to relax and keep their eyes on the center of the monitor.

#### Spatial Working Memory Task and Blink Instructions

The spatial WM span task used in Experiment 2 was identical to Experiment 1 except for: (1) a shorter stimulus presentation (*200 ms* instead of 500 ms) with a total trial duration of 4700 ms, and (2) the color of the cross in the center of the screen for each trial turned from gray to black for 500 ms after stimulus presentation for all trials (Figure 3a). All participants first performed 80 trials with no instructions (“baseline task”). Participants were randomly assigned to the blink group and no blink group in which they performed an additional 80 trials accordingly (“instruction task”; Figure 3b). The participants in the blink group were given the following instructions: “please blink right after the target dots disappear from the screen.” The participants in the no blink group were told to “try your best NOT to blink during the period when the cross is black in color (i.e., first 500 ms of the delay period).” The degree to which the participants followed the instructions were carefully evaluated and calculated as the percentage of trials they followed the instructions (instruction accuracy, blink or no blink during the first 500 ms of the delay period). When analyzing data from the instruction task, outliers based on instruction accuracy (2 SD away from mean) were excluded from final analysis.

**Figure 3.**
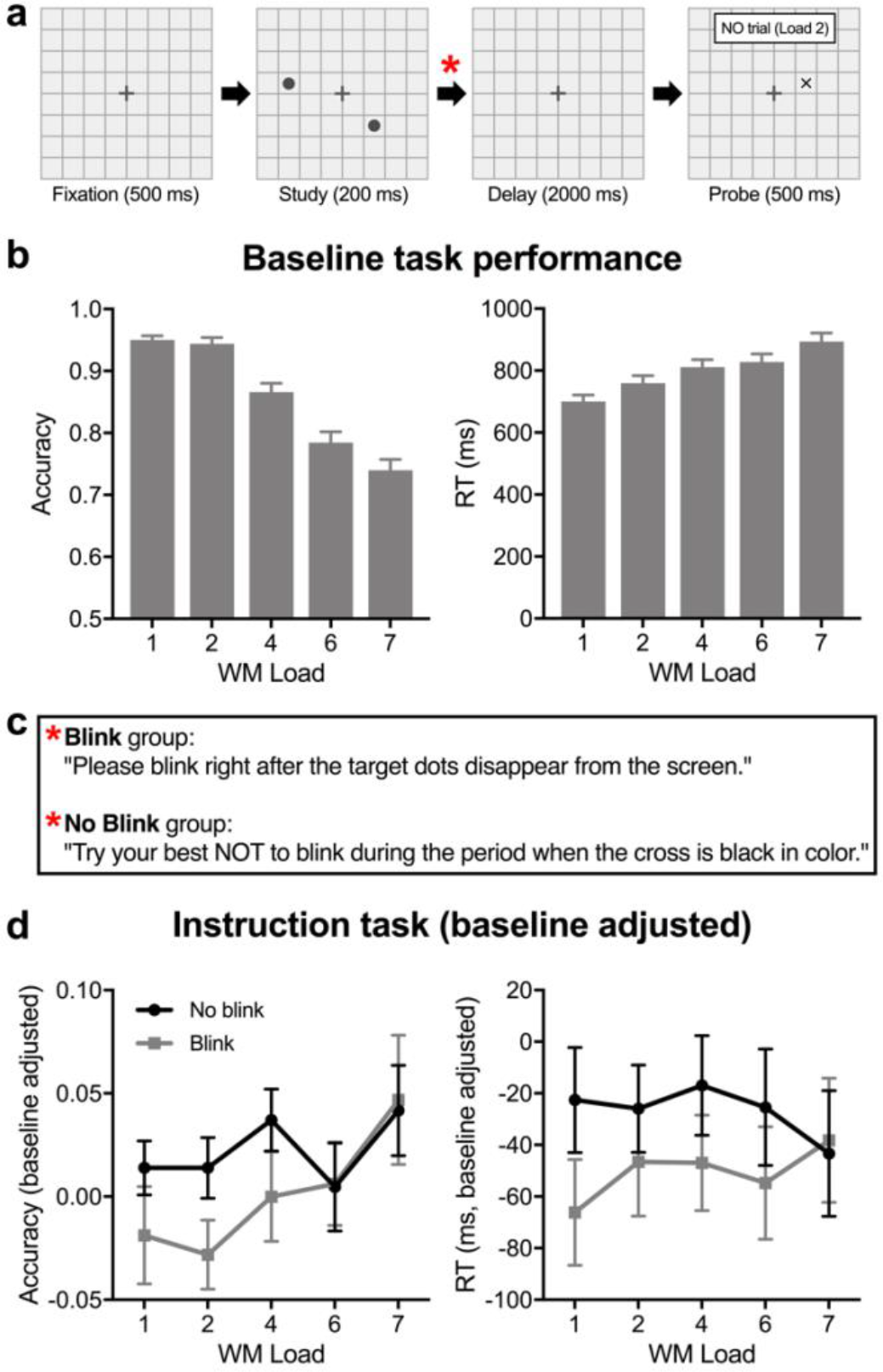
Schematic diagram of spatial WM task and behavioral results for Experiment 2. (a) WM load consisted of one, two, four, six, or seven dots presented in different locations on each trial, followed by a delay. After the delay, a probe appeared prompting a recognition response (yes or no). (b) Baseline task accuracy (left) and RT (right) by WM load. (c) Blink or no blink instructions given to participants. (d) Instruction task accuracy (left) and RT (right) by WM load adjusted using each individual participant’s baseline task performance. Data are expressed as mean ± 1 SEM.

#### Data Analysis

Except for eye blink quantification, the data analysis method and statistical tests were identical as Experiment 1. Number of blinks per trial and time bins were extracted using EyeLink DataViewer (SR Research, Ltd., Ontario, Canada, v1.11.900).

## RESULTS

### Experiment 1

#### Spatial WM Performance

As expected, there was a main effect of WM load for both accuracy (*F*(2.88,170.17) = 97.15, *p* < .001; Figure 1b) and RT (*F*(2.89,170.65) = 80.29 and *p* < .001; Figure 1c). WM capacity or average K-score ranged from 1.29 to 5.67 (SD = 1.27). The participants were grouped into high and low performers using a median split; those with an average K-score larger than the median (3.81) were placed in the high K-score group (*n* = 30, M = 4.89, SD = .50) whereas those below the median were placed the low K-score group (*n* = 30, M = 3.01, SD = .53).

#### Temporal Pattern of Blink Activity During Spatial WM

To examine how blink rate varies over the time course of spatial WM, we calculated SBR using 500-ms time bin. SBR increased after target stimulus offset, decreased over the delay period and increased again after probe presentation and dropped over the ITI (Figure 2a). For correct trials, we conducted a time bin by WM load ANOVA. The main effect of WM load was not significant (*F*(4,2.169) = 1.51, *p* = .22), but the main effect of time bin and WM load by time bin interaction (respectively: *F*(3.34,197.03) = 80.29; *F*(10.89, 642.51) = 3.14; *p’s* < .001) was significant.

As previous studies suggested that blink suppression with increasing cognitive load during stimulus display and subsequent rebound, we examined SBR across the WM load conditions more closely during the stimulus display and the 500 ms after for both the stimulus and probe display (Figure 2a). A *post hoc* ANOVA revealed a main effect of WM load only during the probe display (*F*(2.597,153.211) = 10.87, *p* < .001) with a greater blink suppression towards higher WM load, but not significant for 500 ms after probe display (*F* < 1). A main effect of WM load was not significant during the stimulus display, but trending towards the significant threshold during the first 500 ms of the delay period (*F*(2.438,143.856) = 2.84, *p* = .051). However, there was a significant main effect of performance group (*F*(1,58) = 5.68, *p* = .02), WM load (*F*(2.624,152.170) = 3.14, *p* = .03), and a load by performance group interaction (*F*(2.264,152.170) = 7.207, *p* < .0001; Figure 2b) for this early delay period. With multiple comparisons corrected (Bonferroni), significant differences between the two performance groups were evident at WM load 4 and above. There was also a significant negative correlation across subjects between their SBR during the first 500 ms of the delay period and WM capacity (*r* = −.298, *n* = 60, *p* = .02; Figure 2c). For the first 500 ms after the probe display, there was only a trend towards a significant effect of load by performance group interaction (*F*(3.235,187.633) = 2.48, *p* = .058).

### Experiment 2

#### Spatial WM Performance

In the baseline performance, there was a significant main effect of WM load for both accuracy (*F*(4, 192) = 73.95, *p* < .0001) and RT (*F*(2.836,136.136) = 80.62, *p* < .0001; Figure 3b). The average K-score ranged from 0.5 to 4.96 (SD = 1.14). The participants were again grouped into high and low performers using a median split of their baseline task performance. Those with an averaged K-score larger than the median (3.25) were placed in the high K-score group (*n* = 25, M = 4.13, SD = .53) whereas those below the median were in the low K-score group (*n* = 24, M = 2.30, SD = .79).

Total of forty-seven participants (blink, *n* = 20; no blink, *n* = 27) were included in the final analyses of the instruction task (two subjects from the blink group were removed as an outlier for poor instruction accuracy). We subtracted the accuracy and RT of baseline task performance from instruction task performance for each participant in order to adjust for individual variability at baseline. There were no significant differences between the two instruction groups for both adjusted accuracy (*F*(1,45) = 2.71, *p* = .12, Figure 3d left) and adjusted RT (*F* < 1, Figure 3d right) or load by group interaction in both accuracy and RT (*F*’s<1).

#### Temporal Pattern of Blink Activity During Spatial WM

Similar to Experiment 1, SBR during the baseline WM performance dropped during stimulus and probe presentation but rebounded afterwards (Figure 4a). We conducted a time bin by WM load ANOVA. There was a significant main effect of both WM load (*F*(3.276,157.230) = 6.82, *p* < .001) and time bin (*F*(3.069,147.332) = 26.26, *p* < .001) as well as a significant interaction (*F*(14.427, 692.489) = 2.56 and *p* = .001), indicating SBR changed over time and varied by WM load as in Experiment 1. A *post hoc* ANOVA, however, revealed no significant main effects of WM load for SBR during target display, early delay (first 500 ms), probe display, and 500 ms after probe display (ITI) (all *F*’s<2 and all *p*’s > .3). Unlike Experiment 1, there was no significant WM load by performance group interaction (Figure 4b, 4c; all *F*’s<1) or a correlation between average K-score and SBR (all *r*’s < .2, *n* = 49, *p*’s > .25) in the early delay time bins.

**Figure 4.**
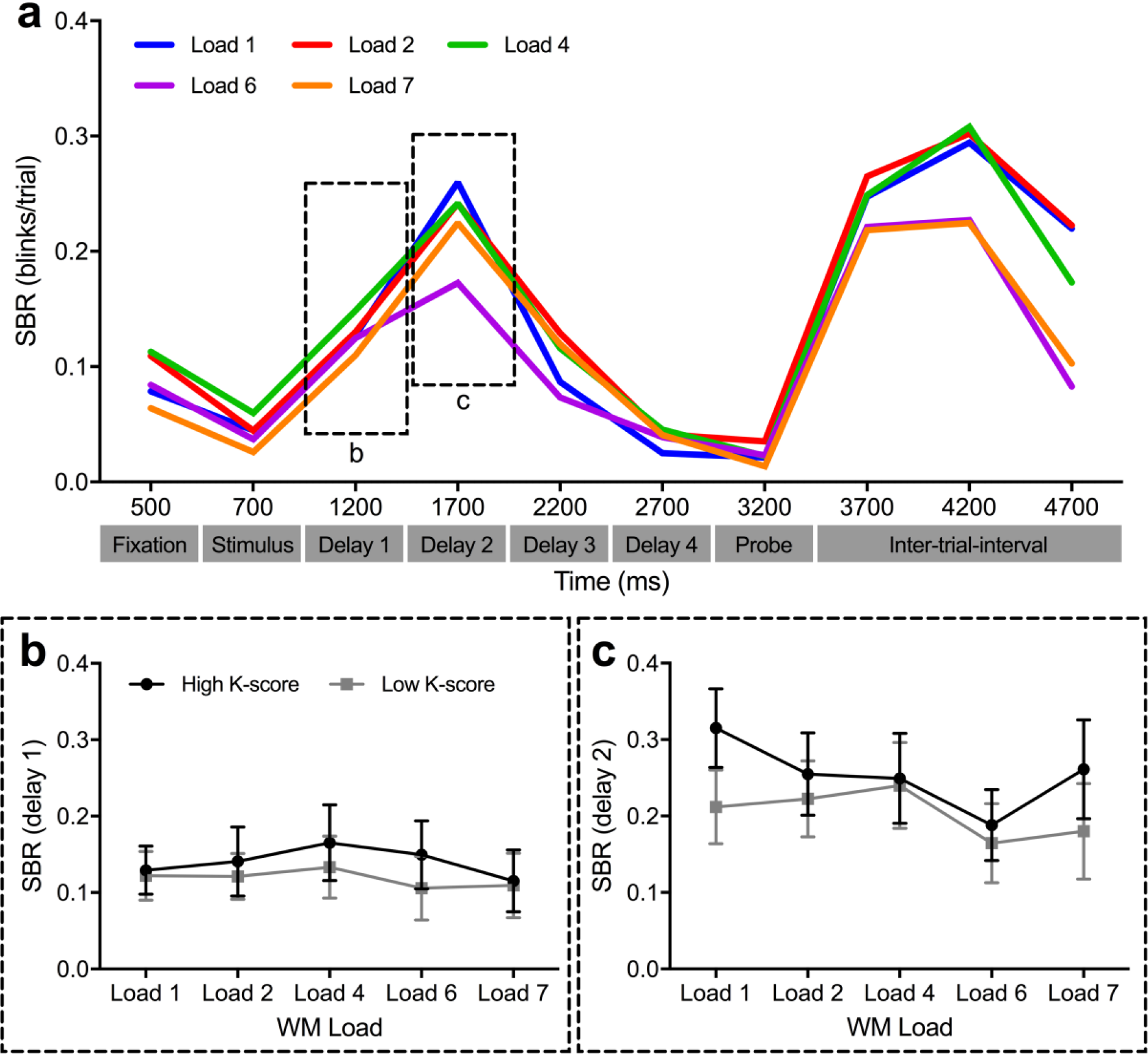
Timecourse of SBR during a full trial and relationship between SBR and WM capacity for Experiment 2. (a) Average SBR across 500 ms time bins for each WM load condition during the baseline task. Time is not graphed to scale on the x-axis (b,c) Differences in SBR between high K-score and low K-score performers by WM load during first 500 ms of the delay period (delay 1, (b); dotted-line box in (a)) and second 500 ms of the delay period (delay 2, (c); dotted-line box in (a)). Data are expressed as mean ± 1 SEM.

#### Spontaneous Eye Blink Activity During Video Watching

SBR while watching the video showed an increase during the first 6 minutes and started to plateau from minute 7 to 10 (Figure 5a), similar to previous results in Kaminer et al. (2011). The average SBR during the 7 to 10 minutes across participants was 20.70 blinks/min (SD = 12.50). There was also a significant positive correlation across subjects between their SBR and WM capacity (*r* = .291, *n* = 48, *p* = .045, one outlier removed; Figure 5b).

**Figure 5.**
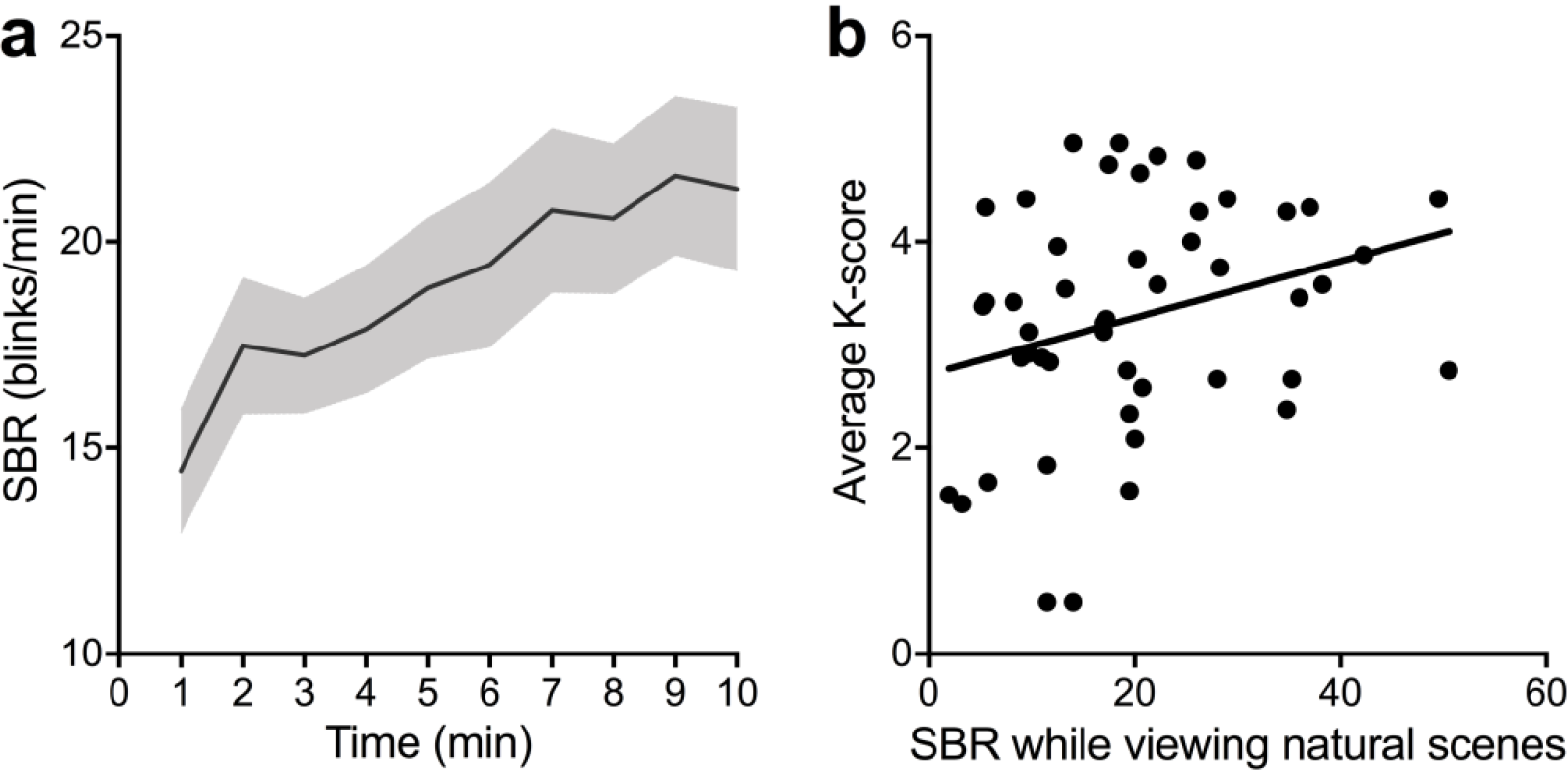
Time course of SBR during video watching and correlation with average K-score. (a) Time course of SBR while watching natural scenes for 10 min. Average SBR for each time bin (1 min each) was calculated and plotted as a function of time. SEM is presented in gray shade area above and under the solid line (mean SBR). (b) Correlation between SBR while viewing natural scenes (average of SBR during 7 to 10 minutes) and average K-score (average of WM load 4, 6, and 7).

## DISCUSSION

### WM event and load dependent temporal changes in SBR

To the best of our knowledge, this is the first study to describe the temporal dynamics in SBR during a spatial WM task with different WM load conditions. In Experiment 1, we showed that SBR is suppressed during stimulus display, early delay and probe display, which are critical for attending to task-relevant visual inputs and information processing (Stern, Walrath, & Goldstein, 1984; Martens, Munneke, Smid, & Johnson, 2006). This suppression was WM load-dependent, with stronger suppression for higher WM loads. These results corroborate previous findings that blink suppression extending beyond the time required for stimulus perception is associated with increased attentional demands and better behavioral performance (Bauer, Goldstein, & Stern, 1987; Fogarty & Stern, 1989; Goldstein, Bauer, & Stern, 1992; Stern & Skelly, 1984; Brookings, Wilson, Swain, 1996; Ohira, 1996; Veltman & Gaillard, 1998; Fournier, Wilson, & Swain, 1999; Van Orden, Jung, & Makeig, 2000; Van Orden, Limbert, Makeig, & Jung, 2001; Ichikawa and Ohira, 2004; Martens, Munneke, Smid, & Johnson, 2006; Siegle, Ichikawa, & Steinhauer, 2008). After suppression of SBR during stimulus and probe display, we also observed a rebound in SBR, which was thought to be compensating for the prior suppression (Stern, Walrath, & Goldstein, 1984). Similar blink suppression and rebound was observed In Experiment 2, though the rebound seemed to continue longer than that in Experiment 1. The shorter stimulus display in Experiment 2 might have led to a stronger suppression of SBR during stimulus display, with the stronger suppression contributed to the delay in SBR rebound. Nevertheless, differences in eye tracking equipment between the two experiments may be inflating this effect. Further studies with parametric manipulation of stimulus duration would be needed to examine potential factors to this difference.

### Individual differences in SBR in association with spatial WM capacity

Here we showed that individual difference in SBR measured during critical events of spatial WM such as the early delay period (Brozoski, Brown, Rosvold, & Goldman, 1979; Sawaguchi & Goldman-Rakic, 1991; Williams & Goldman-Rakic, 1995) is associated with individual spatial WM capacity. In Experiment 1, higher SBR during the early delay period (reflective of WM maintenance-related SBR) was associated with poorer WM capacity. Also, participants with higher K scores showed significantly lower SBR than participants with lower K scores during this period and in a WM load dependent manner. Our finding of this link between successful suppression of SBR during WM maintenance and better WM performance supports the general notion that SBR suppression is relevant for optimal cognitive processing (Stern, Walrath, & Goldstein, 1984). However, we did not observe the same relationship in Experiment 2. The median K-score in Experiment 2 was lower than that in Experiment 1, which may suggest that overall, participants in Experiment 2 were not as successful at suppressing SBR during early delay period compared to participants in Experiment 1. As there were some technical differences in SBR measurements and task parameter differences between the two experiments, these other factors might have also contributed to the discrepancy. Further, sample size may also be another important factor for consideration when studying SBR differences across task events.

The finding of a correlation between SBR during WM maintenance and individual WM performance is also novel. Previous studies examining the relationship between SBR and cognitive task performance have used SBR measured at resting conditions to examine the relationship with task performance (Jongkees and Colzato, 2016). While the direct relationship between individual difference in resting SBR and WM capacity have not been studied, two recent studies have indirectly probed this relationship by studying broader dimensions of executive function (e.g., updating and gating). Higher resting SBR was associated with poorer updating task performance (Zhang et al., 2015) and gating events during a modified n-back task (the reference-back task; Rac-Lubashevsky, Slagter, & Kessler, 2017). While studies did not examine the association between resting SBR and WM maintenance, they suggest that individual differences in resting SBR may be closely related to individual differences in WM performance.

Unlike Zhang et al. (2015), resting SBR was associated with better WM capacity in Experiment 2 in the current study. Two speculations can be made to explain the opposite findings. First, Zhang et al. (2015) used a 3-back task to probe updating, while we used a spatial delayed recognition task to examine WM maintenance. Even within the report by Zhang et al. (2015), they showed that greater resting SBR was associated with better set-shifting and inhibition, suggesting that resting SBR may associate with different dimensions of executive function differently. Second, discrepancies may arise because the relationship between resting SBR and WM performance is, in fact, non-linear. While our results are not sufficient to fully support the non-linearity of resting SBR and WM performance, previous studies showed the possibility of this non-linear relationship. For instance, during a switch task, participants with low resting SBR improved their performance when incentives were at stake, whereas participants with high resting SBR did not (van de Groep, de Haas, Schutte, & Bijleveld, 2017). While the relationship between central DA system and SBR is being questioned (Dang et al., 2017; Sescousse et al., 2018), a study by Cavanagh, Masters, Bath and Frank (2014) showed that cabergoline, unlike other D_2_R agonists (Depue et al., 1994; Ebert et al., 1996; van der Post et al., 2004), increases resting SBR in participants with a low resting SBR and decreases SBR in participants with a high resting SBR. Therefore, when examining the relationship between SBR and cognitive performance, one should not assume a linear relationship and further studies should examine this issue more closely.

### Voluntary eye blinks not associated with spatial WM capacity

As SBR during the early delay period seemed to predict spatial WM capacity In Experiment 1, we expected to see a change in WM performance when blink activity was manipulated through experimental instruction. Contrary to our hypothesis, we did not observe a significant negative impact on WM performance with instructed voluntary blinks during early WM maintenance. Interfering WM maintenance by instructed voluntary blinks has not been widely examined apart from one previous study by Irwin (2014). In Irwin’s study, participants’ K-scores were lower when instructed to blink compared to not blinking during the retention period in a change-detection task. However, their study did not control for individual differences in baseline performance and the instructions were different from our study. In their no blink condition, participants were instructed to not blink throughout the entire stimulus display and retention period (750 ms), while in their blink condition, they were instructed to make a single blink any time during the retention period on the blink trials. Compared to our more controlled no blink condition (limited to early delay), the potentially greater suppression of blinks during the stimulus display in the Irwin (2014) study might have allowed better stimulus encoding and maintenance, which would be consistent with previous studies on blink suppression during stimulus display being associated with better performance (see discussion above). Taken together, voluntary blinks during early spatial WM maintenance do not significantly affect WM performance when taking into account baseline WM performance.

### Limitations

Though SBR during cognitive processing has been studied through multiple approaches, the exact mechanism underlying variation in spontaneous eye blinking is still unknown. This lack of understanding may be due to the complex nature of SBR. Factors such as age, sleep deprivation, usage of contact lenses, etc. (Jonkees & Colzato, 2016) can all change SBR, and some of these factors have not been controlled in our study. Future studies should include an extensive questionnaire to collect information on different factors that influence SBR and control for these factors when examining SBR and cognitive function. Differences in sensitivity of methods measuring eye blinks could also introduce confounds. Future studies should consider multiple eye tracking methods (VEOG and eye tracking camera or high-speed camera). Another limitation in Experiment 2 is that the instruction task was always performed after the baseline task. It is possible that there are practice or fatigue effects, or that the additional cognitive load (i.e., blink instructions) recruited more attention, which might have reduced performance differences between the two conditions in comparison to when no instructions were given. Given the debate on what exact neuromodulatory mechanisms link central DA level and SBR, our findings again raise the possibility that the association between SBR and spatial WM performance may reflect central DA activity. This would need to be confirmed with PET imaging and pharmacological manipulations.

### Summary

Here we used a spatial WM task to examine the temporal patterns of SBR across visual information processing events. We replicated previous findings by showing blink suppression during WM encoding and retrieval, followed by a rebound in blinks. In Experiment 1, we showed that SBR varies in a load-dependent manner such that lower SBR during early WM maintenance was associated with higher WM capacity. These findings indicate that the temporal dynamics of SBR during cognitive tasks is not random, but rather a strategic adaptive behavior for optimal information processing. In Experiment 2, we showed similar results as Experiment 1, but did not see an effect of instructed or suppressed blinks on WM performance. Together, our results suggest that spontaneous, but not voluntary eye blinks may be critically associated with successful spatial WM performance.

## Acknowledgments

We thank Arely Clavel for her technical assistance with collecting EEG data for Experiment 1 and Renee Hartig, Meghan Leonhardt, Anthony Chao for their help with data collection for Experiment 2.

## Notes

1. Note that the SBR suppression during stimuli display in Experiment 2 compared to Experiment 1 was significantly stronger (*F*(1,107) = 30.36, *p* < .0001).
2. The average SBR during the first 500 ms of the delay period in the blink group was 0.91 (SD = 0.09) and in the no blink group was 0.04 (SD = 0.04).
3. To confirm that instructed blinks made by blink group participants were voluntary blinks, we compared average total blink duration in participants in the blink group with an instruction accuracy higher than 0.95 (*n* = 11). Average spontaneous blink and voluntary blink durations were calculated using the early delay period from the “baseline task” trials or “instruction task” trials for each participant, respectively. Indeed spontaneous blinks (M = 161.52 ms, SD = 18.14) were significantly faster than voluntary blinks (M = 298.61 ms, SD = 36.71; *t*(10) = 5.404, *p* < .001 with Wilcoxon signed-rank test). Since the variability in deciding when to raise eyelids during voluntary blinks adds significant time to the total eyelid movement duration, therefore confirming that instructed blinks were indeed voluntary, and not spontaneous eye blinks.
4. In fact, if baseline task performance was not adjusted when analyzing the instruction task performance, there was a significant instruction group by WM load interaction (*F*(1.947,87.609) = 3.26, *p* = .044). *Post hoc* comparison (Bonferroni correction) for each WM load show that the blink group perform significantly better in load 6 and 7 (*t*’s > 2.3 and *p*’s < .01). The instructions may have added some cognitive load and thus heightened the participants’ attention, leading to better performance.

